# SARS-CoV-2 and Mycobacterium tuberculosis co-infection in vitro

**DOI:** 10.1101/2024.08.14.607914

**Authors:** Thays Maria Costa de Lucena, Débora Elienai de Oliveira Miranda, Juliana Vieira de Barros Arcoverde, Mariana Souza Bezerra Cavalcanti, Michelle Christiane da Silva Rabello, Jaqueline de Azevêdo Silva

## Abstract

In less than a year, SARS-CoV-2 (SARS2) has managed to displace *Mycobacterium tuberculosis* (Mtb) as the leading cause of death worldwide due to a single infectious agent. Both pathogens affect the respiratory tract, mainly the lungs. However, the impact that a possible Mtb + SARS2 co-infection can have on the host response is still unknown. Herein we propose (depict) a rigorous system to evaluate the complex interaction between the two infections *in vitro* in a lung epithelial cell line (A549). Overall, the process includes eight steps: (I) Mtb culture, (II) cell maintenance, (III) preparation of viral stocks, (IV) determination of infectious titers, (V) Mtb and SARS2 co-infection, (VI) determination of intracellular bacterial load, (VII) SARS2 viability test, and (VIII) decontamination of supernatants. This comprehensive protocol will allow experimentalists to study the pathogenesis of co-infection *in vitro*, and facilitate collaborative work in the literature.

## INTRODUCTION

*Mycobacterium tuberculosis* (Mtb), the causative agent of Tuberculosis (TB), is responsible for approximately 1.5 million deaths per year^1^. In less than a year, SARS-CoV-2 (SARS2), the pathogen that causes COVID-19, has managed to displace Mtb as the leading cause of death worldwide due to a single infectious agent^2^.

Both pathogens target the human respiratory tract, mainly the lungs^3^. Several case reports of Mtb + SARS2 co-infection have been demonstrated in clinical settings^4–6^, suggesting a possible bidirectional interaction between them, in which the immunosuppression induced by SARS2 infection may contribute to the progression of TB from a latent to an active disease state. In turn, the temporary immunosuppression induced by TB may increase the susceptibility for complications inpatients to COVID-19^7,8^.

Studying both pathogens in a laboratory setting requires prior isolation of the bacteria and viruses individually and their propagation in tissue culture. We developed a system using Mtb and SARS2 strains that were separately cultured and subsequently plated, in mono-infection and co-infection, on a lung epithelial tissue cell line (Fig. 1), the A549 cell. Here, we present a protocol that prioritizes the essential aspects of cell line maintenance and provides a detailed method for infection by pathogens that are structurally and biologically distinct, as well as performing the viability of mono-culture and co-culture, without cross-contamination. In addition, we demonstrate how to obtain decontaminated RNA and protein samples from these cultures, and how they can be transported from a biosafety level (BSL) 3 (BSL-3) laboratory to BSL-2. Ultimately, this in *vitro* Mtb and SARS2 co-infection system may provide the ability to explore how potential pathways may be activated and establish new insights into the concomitant pathogenicity of these diseases.

**Figure 1.**
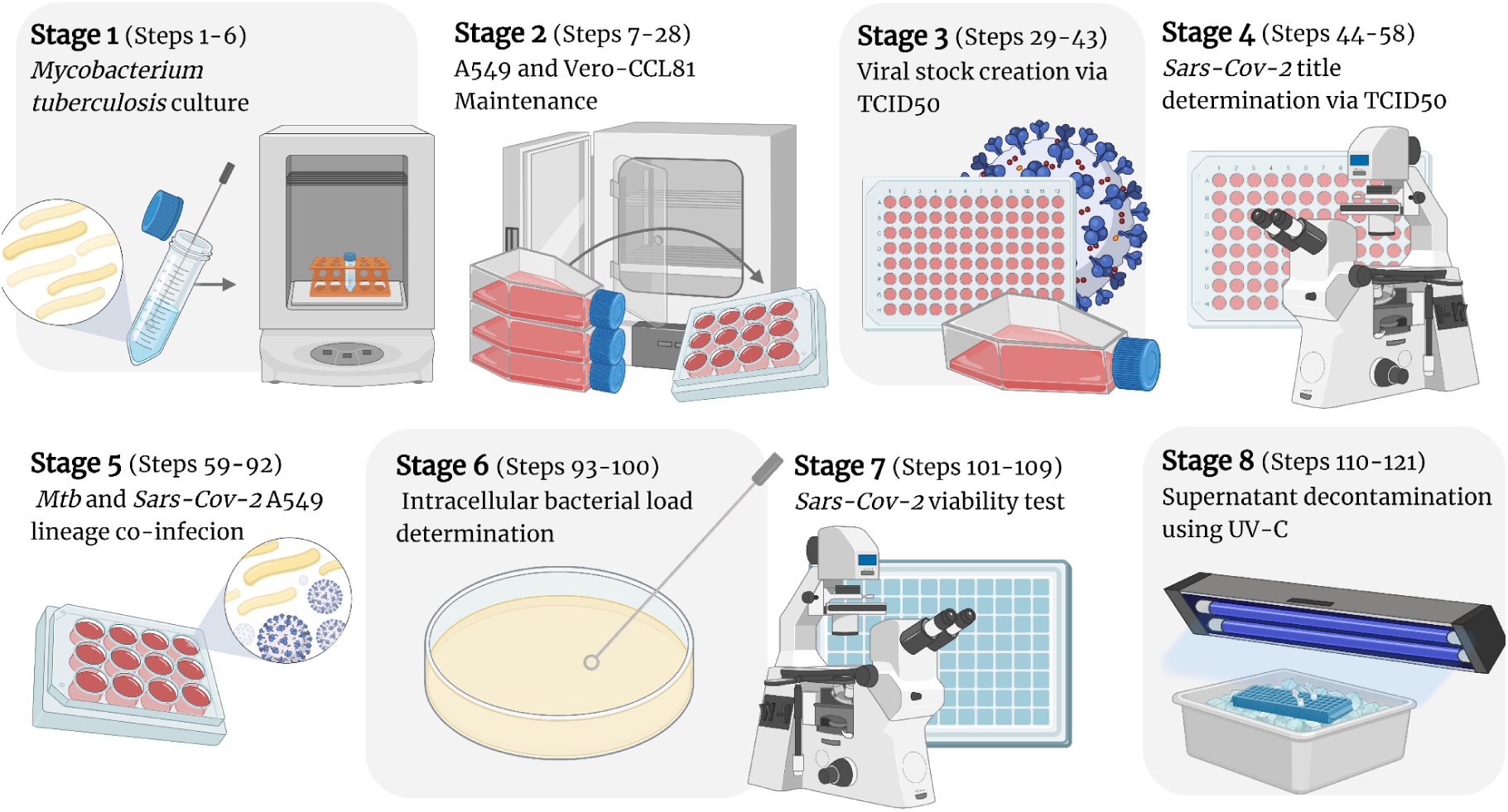
Workflow for Mtb and SARS2 co-infection.

### Protocol Development

The emergence of the COVID-19 pandemic has led researchers around the world to join forces in order to understand the rapid spread of SARS2 which, in turn, neglect the attention to other infectious diseases such as TB, resulting in an increased number of deaths^9^. Additionally, a drastic impact on diagnoses notification raised questions about whether, in TB-endemic areas, underreporting of possible co-infections (Mtb + SARS2) could influence the number of serious cases of COVID-19^1^.

In response, our research group developed an Mtb + SARS2 co-infection system that allows evaluating the viability of the system in mono-infection and co-infection, and obtaining cell extracts, such as RNA and proteins, to study the influence of bacterial and/or viral infection, and potential response pathways that may be activated. Briefly, three independent experiments were performed to demonstrate the replicability of the system. In a 12-well culture plate, the groups were distributed as follows: (1) A549, (2) A549 + Mtb, (3) A549 + Mtb + SARS2, and (4) A549 + SARS2, all were arranged in biological triplicate. After the incubation time, cell extracts were obtained for: (1) evaluation of the intracellular bacterial load, using 7H10 medium plates and direct counting of colony-forming units (CFUs), (2) viral viability test using the TCID50 method, analyzing the cytopathic effect (CPE) by optical microscopy and subsequent obtaining of plaque-forming units (PFUs), (3) Viral and/or bacterial RNA for subsequent gene expression assays, (4) and whole viral and/or bacterial protein obtained from the supernatant for subsequent analysis by flow cytometry. More detailed information about using this method, including troubleshooting information, is provided in this protocol article.

### Protocol Overview

Stage 1 of the protocol (steps 1-6) refers to the cultivation of Mtb. Due to the slow growth of this bacterium, this stage must occur before the others. Mtb presents an ideal concentration of bacteria in the culture for approximately 28 days, during which stages 2, 3 and 4 can be performed. Stage 2 (steps 7-28) corresponds to the thawing of the host cells, maintaining them so that satisfactory cell growth occurs, confluence ≥ 90% is obtained and infections continue.

Stages 3 (steps 29-43) and 4 (steps 44-58) involve the preparation of viral stocks and determination of infectious titers by TCID_50_, respectively. This last stage shows the experimentalists whether there was a CPE, and the quantity of viral particles present in this same culture, this value being subsequently represented by TCID50/ml, and finally used as a control to verify whether there was an increase or decrease in viral activity in the culture infected by SARS2. After 28 days of Mtb growth with certified optical density (OD) at 0.6 (value representing the ideal concentration of bacteria), confluent host cells and determined infectious titers, stage 5 (steps 59-92) can be performed to co-infect the host cells with both pathogens. The culture plate configuration should be presented in biological triplicate, with wells with only A549 cells, wells mono-infected with Mtb, mono-infected with SARS2 and co-infected with Mtb and SARS2. At this same stage, aliquots of the cultures are made and sent for the cell extracts, RNA and host protein, intracellular bacterial load test (stage 6, steps 93-100) and SARS2 viability test (stage 7, steps 101-108). We also present an optional stage (Stage 8, step 109-121) for the decontamination of supernatants obtained from cell culture. Stages 3 to 8 must be performed exclusively in BSL-3.

### Comparison with other methods

The first protocol ever published on co-infection between two distinct pathogens reports that bacterial infection, specifically *Staphylococcus aureus*, is one of the most common complications of COVID-19^10^. However, with this protocol we aim to evaluate how Mtb can previously influence subsequent infection by SARS2, and how this co-infection can modulate the response of infected host cells. It is possible to quantify the kinetics of viral and bacterial replication through two methods: (1) RNA extraction, using Trizol, with disruption of the nuclear membrane, and (2) host proteins, evaluating cellular activity by flow cytometry, requiring that the plasma membrane of these cells be intact. For the late, a decontamination step using ultraviolet radiation type C (UV-C) light is necessary, where there is no protein degradation. Here we demonstrate how to enable decontamination using this methodology in the laboratory, and how to transfer these samples from BSL-3 to BSL-2, ensuring safety for the experimentalists.

Vero-E6 cells are widely used for SARS2 infection, due to their greater expression of the ACE2 receptor and ease handling and support for high levels of viral replication. Tests with Vero-CCL81 showed similar capacity in viral propagation^11^. Due to its availability in our laboratories, we demonstrate here that Vero CCL81 can be used as an efficient alternative for SARS2 infection.

### Expertise required to implement the method

Implementation of this protocol requires expertise in mammalian tissue culture techniques, as well as experience in handling highly pathogenic microorganisms. All experiments with Mtb and SARS2 are performed in a BSL-3 laboratory. It is thus incumbent upon those engaged in experimental work with these pathogens to receive the requisite training and certification for access to such high-containment facilities.

### Limitations

Our experimental project is to co-infect a lung epithelial tissue cell line with *Mtb* and SARS2, and obtain samples to verify the immune response pathways that may be activated in the face of co-infection. This project has four limiting points: (I) Mtb is a slow-growing bacterium, which makes it difficult to obtain faster results, taking approximately ∼60 days to obtain culture results, considering thawing of bacterial strain until definition of the intracellular bacterial load. (II) The viral stocks produced and the samples must be frozen, and they cannot be thawing-refreezing, as this can reduce the sample titer. (III) the BSL-3 used for this experiment has a facility with essential equipment for handling the pathogens. The execution of other experiments with cell extracts from these cultures, such as RNA and protein, requires an additional step using reagents such as Trizol and UV-C, respectively, to ensure proper decontamination and no infectivity upon leaving BSL-3. Due to the wide variety of equipment with UV-C light, specific analyses were performed with a radiometer to determine the adequate incidence of light, as well as the necessary time of this radiation. To ensure decontamination efficiency, another culture was performed in biological triplicate. (IV). The host cells must be suitable for continuous replication of both pathogens: Mtb and SARS2. For this, the A549 cell line was selected due to the presence of cell surface receptors, which allow the entry of Mtb, in addition to expressing the angiotensin-converting enzyme 2 (ACE2) receptor, being suitable for the entry and replication of SARS2.

## MATERIALS

### Biological Materials

#### Cells

- Vero (CCL81)
- A549 (CCL185)

**Critical:** it is imperative to ensure that cell viability and harvest densities remain stable and that the cells utilized in this protocol are free from contamination with mycoplasma.

#### Virus

- Samples of virus strains can be acquired through national/international biorepositories or academic sources. The SARS2 strain used in this experiment was hCoV-19/Brazil/PE-FIOCRUZ-IAM4372/2021.

#### Bacteria

- Bacterial samples can be acquired through national/international biorepositories or academic sources. The bacteria used in this experiment was Mtb H37Rv (UFPEDA 82).

**Critical:** all procedures with SARS2 and Mtb must be performed in a BSL-3 environment using standard BSL-3 practices.

### Reagents

#### Cell Culture

- Phosphate Buffer Saline (PBS) (GIBCO, cat. 10010023)
- Dulbecco’s Modified Eagle Medium (DMEM) with 4.5g/liter of D-glucose, L-glutamine, with sodium pyruvate (Capricorn Scientifc, cat-no. DMEM-HPA)
- Fetal Bovine Serum (FBS) (Gibco, cat. 12657029)
- Penicilin-Streptomycin (10.000U/mL) (Gibco, cat. 15140122)
- Trypsin-EDTA (Sigma-Aldrich, cat. T4049)
- Trypan Blue (Gibco, cat. 15250061)
- Cell-counting chamber slides with trypan blue stain (ThermoFisher Scientific, cat. no. C10312)

#### Infection

- Ogawa-Kudoh BK (Laborclin, ref. 510156)
- Middlebrook 7H9 medium (BD DIFCO, cat. DF0713-17-9)
- Middlebrook 7H10 medium (BD DIFCO, cat. 11799042)
- Oleic Acid-Albumin-Dextrose-Catalase (OADC) (BD BBL, cat. 211886)
- Tween 80 (ThermoFisher Scientific, cat. 28329)
- UltraPure™ Glycerol (ThermoFisher Scientific, cat. 15514011)

### Equipment

- -80 °C Freezer
- Class II Biological Safety Cabinet (ControlAir Biosafe)
- Class III Biological Safety Cabinet (ControlAir Biosafe)
- CO_2_ incubator (Heraeus HERAcell incubator, Thermo Fisher)
- Bain-marie (SolidStell, ref. SSDC15L)
- Cell counter
- Shaker Incubator (e.g., Benchmark INCU-SHAKER™)
- Inverted microscope (e.g., Nikon Eclipse Ts2R)
- Light microscope (e.g., Nikon Eclipse Ts2-FL)
- Microcentrifuge (Eppendorf™, cat no. 5429000015)
- Refrigerated Benchtop Centrifuge (Eppendorf™, cat. no 05-400-61)
- 6-well cell culture plate (Fisher Scientific, cat. no. 07-200-83)
- 12-well cell culture plate (Fisher Scientific, cat. no. 07-200-82)
- 96-well cell culture plate (Eppendorf, cat. no. 951020320)
- T25 Cell Culture Flask (Thermo Scientific, cat n. 156340)
- T75 Cell Culture Flask (Thermo Scientific, cat n. 156472)
- Newbauer Chamber
- Sterile syringes, 1 ml (e.g., Fisherbrand, cat. no. 14-955-464)
- Sterile syringes, 5 ml (e.g., Fisherbrand, cat. no. 14-955-458)
- Conical centrifuge tube, 15 ml (Thermo Scientific, cat. no. 339650)
- Conical centrifuge tube, 50 ml (Thermo Scientific, cat. no. 339653)
- Microcentrifuge tubes, 1.5 ml (Thermo Fisher Scientific, cat. n.° AM12300)
- Tubos com tampa de rosca O-ring, 2 ml (Sarstedt Inc, cat. n° 72.694.006)
- Filtros Corning SFCA 0,22 µm (Fisher Scientific, cat. n° 09-754-13)
- Erlenmeyer (e.g., Fisherbrand 500 ml; Fisher Scientific, cat. no. FB 501500)
- Serological pipette, 5 ml (Fisher Scientific, cat. n.° 501214573)
- Serological pipette, 10 ml (Fisher Scientific, cat. n.° 501214572)
- Serological pipette, 25 ml (Fisher Scientific, cat. n.° 501214571)
- Micropipettes (Kasvi)
- Micropipette tips, sterile, filtered, 10, 20, 200 e 1.000 µl (Fisherbrand or equivalent)
- Petri dishes 100 x 15 mm (Thermo Scientific, cat. no. 150350)
- Parafilm (Fisher Scientific, cat. n° NC9595547)

### Reagent Setup

#### Maintenance medium

The maintenance medium contains components that are necessary for cell proliferation and culture adaptation after thawing and passages. It can be prepared in a BSL-1 or BSL-2 laboratory in a biosafety cabinet. In a 500 ml bottle of DMEM modified to contain 4 mM L-glutamine, 4.500 mg/L glucose, 1 mM sodium pyruvate, add 5 ml of penicillin-streptomycin (100×; final concentration: penicillin, 100 IU; streptomycin, 100 µg/ml). Since it is a maintenance medium, heat-inactivated FBS at 10% (vol/vol) should be used; add 50 ml of this component to the bottle of DMEM previously supplemented with antibiotics. Mix the bottle well with circular movements or by inversion. Distribute 50 ml aliquots into 50 ml conical tubes and store at 4°C for ≤ 6 weeks.

#### Growth medium for Mtb – 7H9

Middlebrook 7H9 medium is a liquid culture medium specific for mycobacteria. It can be prepared in BSL-1 or BSL-2 laboratories. Weigh 2.35 g of MiddleBrook 7H9 Medium and place it in a sterilized 500 ml bottle. Add 449 mL of ultrapure water and 1 mL of glycerol. Perform circular movements to homogenize. Autoclave at 121°C for 10 minutes.

To continue with the supplementation, the medium must be at room temperature. Send the bottle to the BSL-3 laboratory and perform the distribution of 45 mL of the medium in 50 mL conical tubes in the biosafety cabinet, and add the equivalent of 10% OADC (5 mL) and 0.05% Tween 80 (25 µL) to each tube. Mix the contents by vortexing the tube for 10 s or until the Tween 80 is completely dissolved. Store at 4°C for ≤ 6 weeks.

#### Growth medium for Mtb – 7H10

Middlebrook 7H10 medium is a solid culture medium used for the cultivation of mycobacteria. Prepare the 7H10 medium in a BSL-2 laboratory. Weigh 4.75 g of MiddleBrook 7H10 Medium and transfer the contents to a 500 mL Erlenmeyer flask. Add 223.75 mL of ultrapure water and 1.25 mL of glycerol, microwave for 3 minutes, and every 1 minute remove and perform circular movements to more efficiently dissolve the contents. Autoclave at 121°C for 10 minutes.

After sterilization, this medium must be supplemented. Wait for it to cool to room temperature and use a thermometer to check the temperature of the medium, still in a liquid state. When it reaches 40°C, add 25 ml of OADC to the Erlenmeyer flask and quickly and safely plate by inserting 25 ml of supplemented medium into each disposable Petri dish. Allow the medium to dry completely and seal the plates with parafilm. Place a control plate in the incubator at 37°C for 24 h to check for possible contamination. If there is no contaminant growth, transport the plates to the BSL-3 laboratory and store at 4°C for ≤ 2 weeks.

#### Lysis Buffer Stock Solution

Prepare a lysis buffer in a BSL-2 laboratory. In a 250 mL vial, add 0.1% v/v Tween 80 in 250 mL deionized water. Autoclave the material at 121°C for 10 min, allow it to cool to room temperature, and filter the entire volume through a 0.22µM filter. Transport to BSL-3 and store at room temperature for ≤6 weeks.

#### Dilution Buffer Stock Solution

Prepare BSL-2 laboratory dilution buffer. In a 250 mL bottle, add 0.05% v/v Tween 80 in 250 mL PBS. Autoclave the material at 121°C for 10 min, allow it to cool to room temperature, and filter the entire volume through a 0.22µM filter. Transport to BSL-3 and store at room temperature for ≤6 weeks.

#### Virus and Bacterial Infection Medium

The infection medium is adapted to promote infection of cells, specifically viruses and bacteria. Prepare the viral and bacterial infection medium in a BSL-1 or BSL-2 laboratory. To prepare an infection medium, 2% FBS is required. Add 10 ml of heat-inactivated FBS, 5 ml of penicillin-streptomycin (100×; final concentration: penicillin, 100 UI; streptomycin, 100 µg/ml) to a 500 ml bottle of DMEM modified to contain 4 mM L-glutamine, 4500 mg/L glucose, 1 mM sodium pyruvate. Mix the bottle thoroughly by swirling or inverting. Distribute 50 ml aliquots into 50 ml conical tubes. Transport to the BSL-3 laboratory and store at 4°C for ≤ 6 weeks.

## PROCEDURES

### Stage 1: Mycobacterium tuberculosis culture

**Timing:** up to 28 d.

**Critical Step:** this procedure must be performed exclusively in a BSL-3 laboratory.

**Day 1:** Preparation of bacterial culture

**Timing:** 1 h inoculation, 4 d incubation.

1. Add 5 mL of Middlebrook 7H9 Medium (supplemented with 10% OADC and 0.05% Tween 80) to a 50 mL conical tube.
2. Using a disposable bacteriological inoculation loop, scrape the Mtb into Ogawa-Kudoh medium (Fig. 2), ensuring that there are enough bacteria on the loop, carefully transfer it to the tube with 7H9 medium and mix the bacteria into the medium.
3. Place the culture obtained in the previous step in the shaker incubator at 37°C for 4 days.

**Figure 2.**
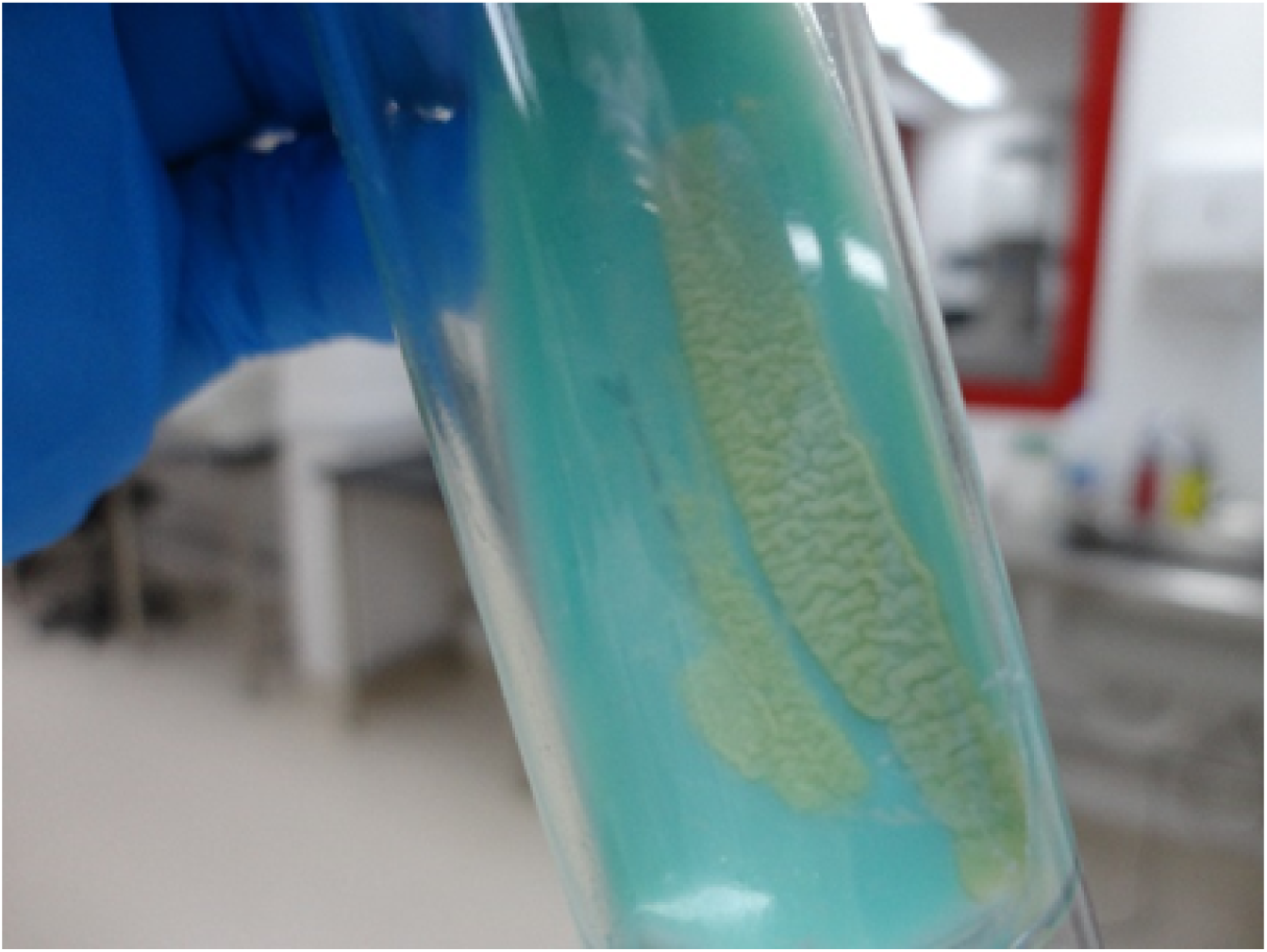
Mtb H37Rv strain (UFPEDA82) in Ogawa-Kudoh medium.

**Dia 4:** Maintenance of bacterial culture

**Timing:** 1 h for culture maintenance

4. Remove the culture from the incubator, and in the biological safety cabinet, add 7 ml of 7H9 medium to the bacterial culture to replenish nutrients, maintain pH and reduce cell density, thus achieving efficient growth.
5. Return the culture to the shaker incubator at 37°C.
6. Observe weekly until reaching 28 days of cultivation.

**Critical Step:** Mtb is a slow-growing bacterium, taking about 28 days to reach the ideal logarithmic phase. If the bacterial culture shows excessive turbidity before 28 days, check the optical density (OD) on the spectrophotometer. More details on how to perform the measurement are in stage 5, step 62.

### Stage 2: cell maintenance (A549 and Vero-CCL81)

**Timing:** 10 d for thawing and confluence of cells

**Critical:** the procedures at this stage can be performed in BSL-1 or BSL-2 laboratories.

**Day 1:** thawing, replating and cell counting

**Timing:** 1 h for thawing and inoculation, 2 – 3 d for growth and 90% confluence

7. Pre-heat the flask containing cell maintenance medium to 37°C in a water bath.
8. Remove the cryovial containing the cells from the liquid nitrogen and thaw them at 37°C in a water bath.
9. In a biosafety cabinet, carefully transfer the cells with p1000 to a 15 mL conical tube and add 9 mL of the pre-warmed maintenance medium prepared in step 7.
10. Centrifuge the cells at 1,200 RPM for 5 minutes at room temperature. This process removes any possible freezing residue and forms a pellet.
11. Discard the supernatant.
12. Resuspend cells using p1000 in 1 mL of pre-warmed maintenance medium, mix gently by pipetting and transfer to a 15 mL conical tube.
13. Count viable cells using a cell counter. In a 1.5 ml eppendorf tube, mix 10 µl of the cell suspension with 90 µl of Trypan blue. Add 10 µl of this mixture to one side of a slide inserted into the Newbauer Chamber and count the viable cells in the four quadrants.
14. The cells must have a density of 1 × 10^5^ live cells/cm^2^.

**Critical:** it is recommended that cells be seeded at a high density to facilitate a more rapid recovery following the thawing process.

15. Transfer the volume of cells for seeding obtained in step 14 to the flask and calculate the difference in maintenance medium required.
16. Pipette the pre-warmed maintenance medium into a desired size bottle (T25 bottle should total 5 ml and T75 bottle should total 15 ml).
17. Place the flasks with the cells to grow in a CO_2_ incubator (5%) at 37°C until reaching 90% confluence. Check the cultures daily.

**Critical:** The cells should reach a confluence of 90% and be prepared for division after a period of 2–3 days. Prior to dividing the cells, a visual inspection should be conducted under a light microscope to ascertain whether recovery from freezing has occurred. The cells should be attached to the vial with only a few cells floating around, and their morphology should be unaltered. In the event that cells have not recovered from the thawing process, they often exhibit a more rounded morphology and a reduced degree of connection to neighboring cells. In the event that the cells fail to recover from the freezing process, it is advisable to begin anew with a fresh vial of cells.

**Critical:** A549 cells grow in clusters, so it is recommended to use T25 flasks for their cultivation.

**Days 3-4:** cell division and replication

**Timing:** 1 h for division, 4-6 d for two passes

**Critical:** This section delineates the methodology for two passes, which is imperative to guarantee recuperation from freezing and a stable growth rate prior to initiating subsequent experiments. The process may take up to ten days from the time of thawing.

18. In a water bath at 37°C, pre-heat the maintenance medium, trypsin-EDTA and PBS.
19. Aspirate the medium contained in the flasks with a pasteur pipette or automated serological pipette and wash the cells once with PBS. Aspirate the PBS and add trypsin-EDTA to cover the monolayer. Incubate for 5 minutes in an incubator at 37°C.

**Critical:** For T25 and T75 flasks, add 3 ml and 9 ml of PBS, respectively. For trypsin-EDTA, add 2 ml for T25 flasks and 6 ml for T75 flasks.

20. Check the detachment of cells under an optical microscope (Fig. 3).

**Figure 3.**
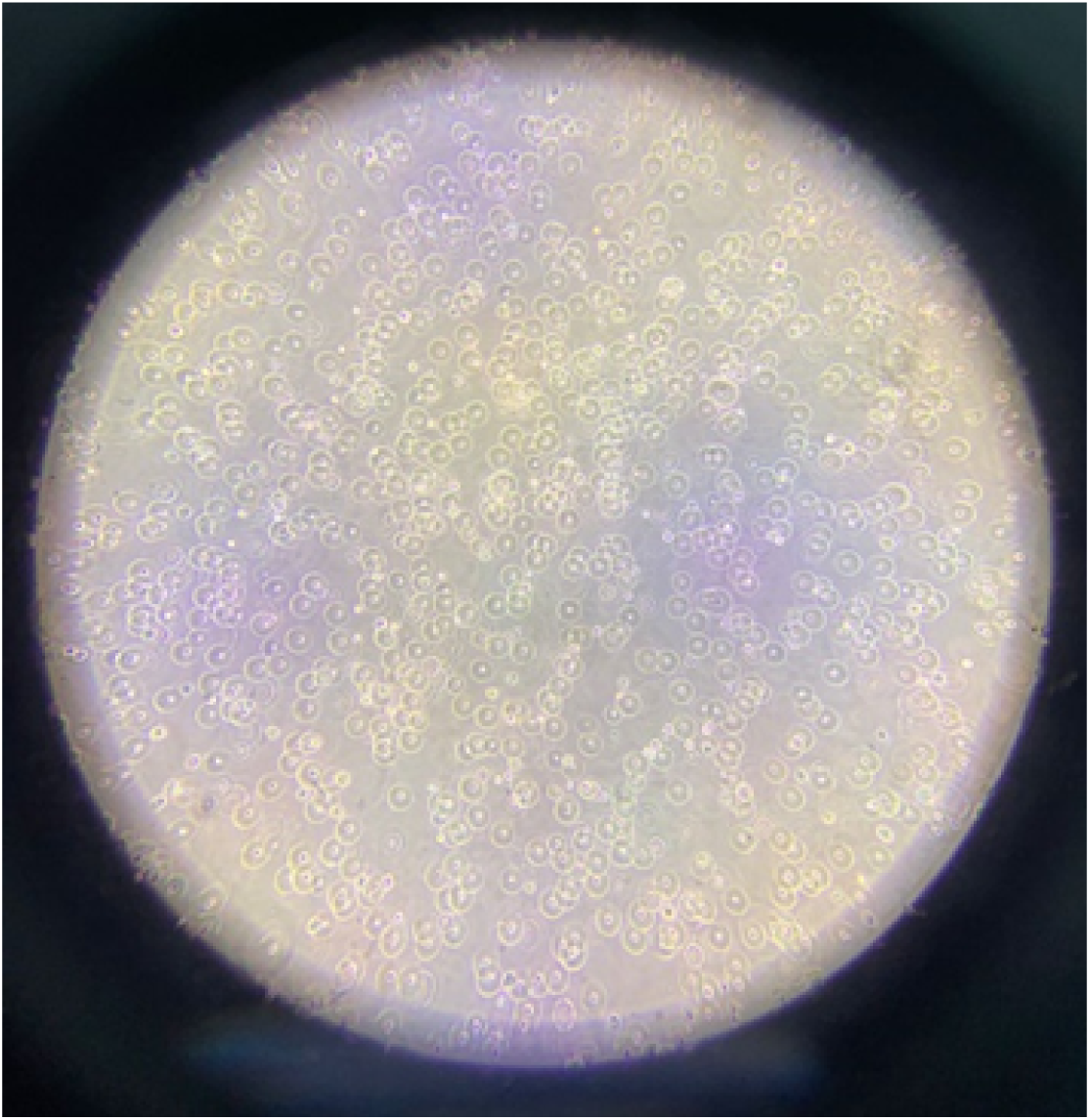
A549 cells in the trypsinization process.

**Critical:** the A549 cell line forms clumps during growth. Under the microscope, it is possible to observe the cells detached from the bottom of the vial, but they are still aggregated. Tap the bottle gently to break up these clumps into single cells.

21. Transfer the cells to a 15 or 50 ml conical tube and stop trypsinization by adding a pre-warmed maintenance medium in the same volume of trypsin-EDTA as in step 19, considering the size of the flask.
22. Pellet cells at 1,200 RPM for 5 min at room temperature.
23. Discard the supernatant.
24. Resuspend cells in 1 ml of maintenance medium.
25. Count live cells as per step 13.
26. Transfer the required volume of cells and maintenance medium for seeding into the desired flask.

**Critical:** for 90% confluence in 2 days, seed cells at 4 × 10^4^ live cells/cm^2^. For the same confluence in 3 days, seed at 2.5 × 10^4^ live cells/cm^2^.

27. Place the flask in an incubator at 37°C with 5% CO_2_. Check the cultures daily.
28. Repeat steps 18 through 28 once before using the cells for experiments.

### Stage 3: preparation of viral stocks by TCID_50_

**Timing:** 5 d

**Day 1:** Vero-CCL81 cell plating

**Timing:** 1 h and overnight incubation

29. In a BSL-1 or BSL-2 laboratory prepare 1 vial of T25 and 1 vial of T75 with Vero-CCL81 cells at a density of 1 × 10^4^ cells/cm^2^.
30. Place the flasks in an incubator at 37°C with 5% CO_2_ overnight.

**Day 2-5:** inoculation of Vero-CCL81 cells with SARS2 strain

**Timing:** 2 h for inoculation and 3 d for incubation

**Critical:** procedure with SARS2 should be performed in a BSL-3 laboratory.

31. Transfer the flasks with Vero-CCL81 cells to the BSL-3 laboratory.
32. Remove the cell maintenance medium, and then:

a) T25 (control): add 400 µl of infection medium and mix carefully with circular movements.
b) T75 (infection): add 1 ml of the viral strain aliquot and mix carefully with circular movements.
33. Place the flasks in an incubator at 37°C with 5% CO_2_ for 1 hour.

**Critical:** every 15 minutes, observe the bottles and make circular movements to prevent the monolayer from drying out.

34. Using p1000, add 4.6 ml of infection medium to the T25 bottle and 14 ml of infection medium to the T75 bottle.
35. Store the vials in the incubator at 37°C with 5% CO_2_ for 72 h.
36. Remove from the incubator and store the bottles in the freezer at -80°C for 24 h.

**Day 6:** virus harvesting

**Timing:** 2 h

37. Thaw the flasks at room temperature.
38. View the flasks under an optical microscope. Observe whether there is contamination in vial T25 and whether the CPE is visible in vial T75 (Fig. 4).
39. Discard the T25 flask.
40. Transfer the contents of the T75 flask to a 50 ml conical tube.
41. Remove cellular debris by centrifugation at 4,500 RPM for 10 min at 4°C.
42. Transfer the supernatant to a new 50 ml conical tube and discard the tube containing the pellet.
43. Identify tubes with screw caps and sealing rings and make aliquots of 150 µl each to be used in the next step.

**Figure 4.**
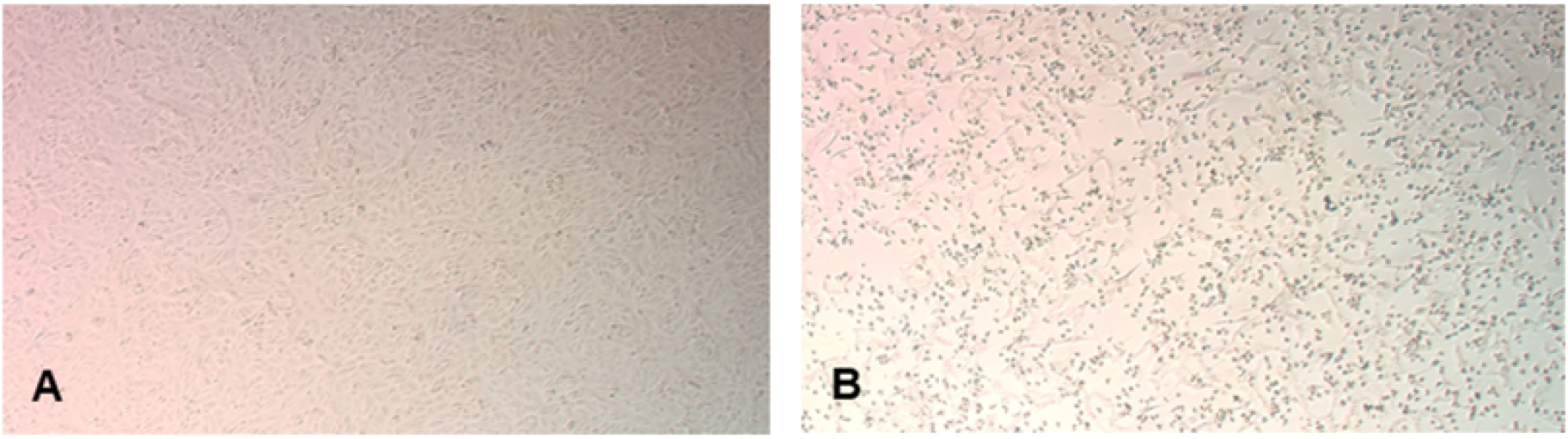
Vero CCL-81 cells (control and SARS2 infected). In A) Uninfected cells used as controls; B) SARS2 infected cells.

### Stage 4: determination of SARS2 infectious titers by TCID_50_ assay

**Timing:** 5 d

**Day 1:** plating of Vero-CCL81 cells for infection

**Timing:** 1 h for infection and incubation overnight

44. Prepare Vero-CCL81 cells in two 96-well plates at 1 × 10^4^ cells/well (Fig. 5a).

**Figure 5.**
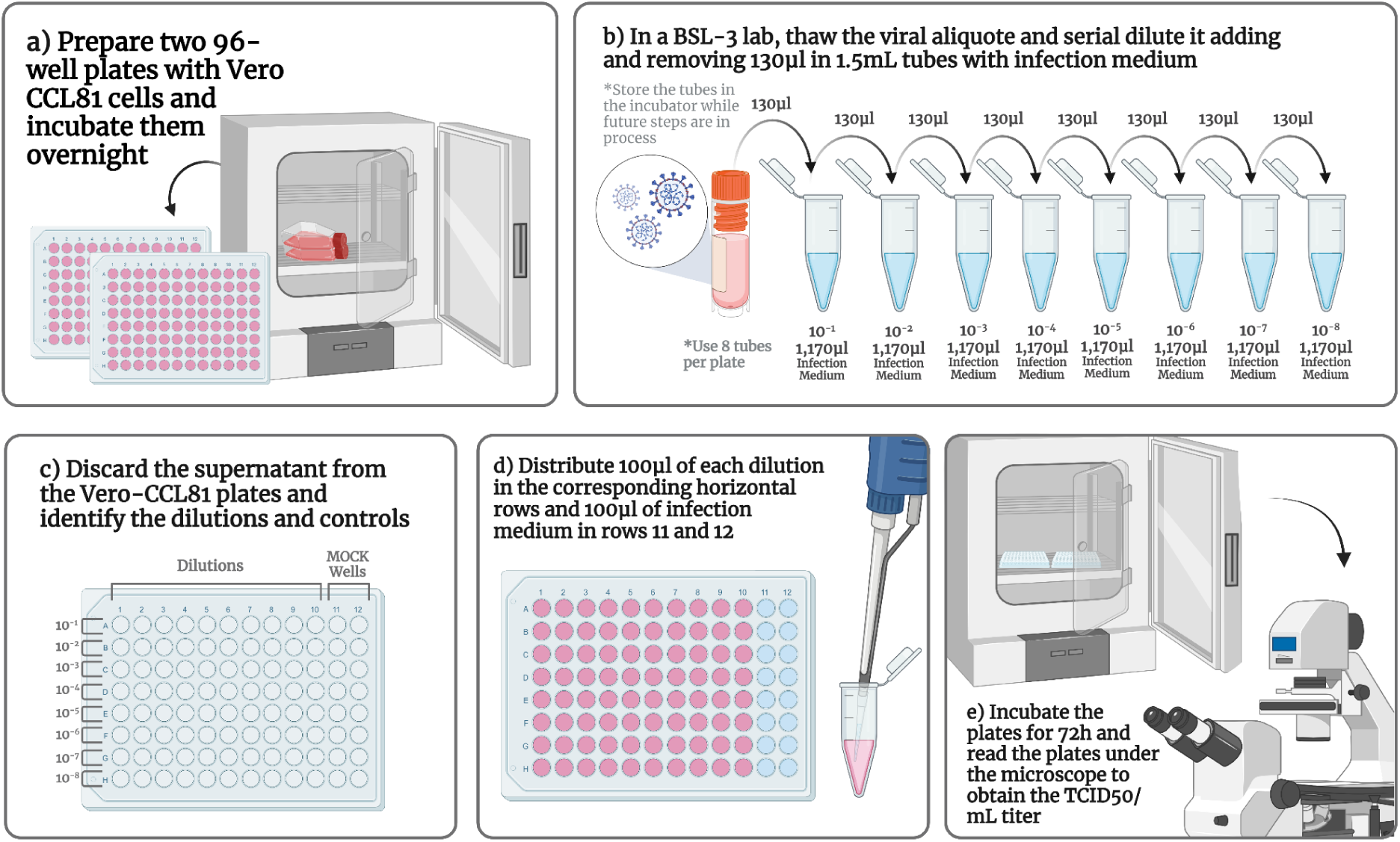
Workflow for TCID_50_ assay. a) Vero-CCL81 incubation. b) Dilution of the viral aliquot. c) Identification of dilutions and MOCK in the 96-well plate. d) Insert the dilutions into the corresponding wells. e) Incubate the plates.

**Critical:** the preparation of two plates is to demonstrate experimental replication, ensuring experimental replication, consistency of results in different replicates, reduction of experimental errors, such as contamination and inadequate handling, and robust statistical analysis.

45. Place the plates in a 37 °C incubator with 5% CO_2_ overnight.

**Critical:** After the incubation period, check under the microscope whether the cells have formed a compact confluent monolayer in each well.

**Critical:** This stage can be performed in a BSL-1 or BSL-2 laboratory. The remaining stages must occur under BSL-3 containment.

**Day 2:** TCID_50_ assay

**Timing:** 2,5 h for inoculation, 72 h for incubation

**Critical:** this procedure with infectious SARS2 must be performed in a BSL-3 laboratory.

46. Transfer the plates to the BSL-3 laboratory and place them in a 37°C incubator with 5% CO_2_ until virus inoculation.
47. Thaw a viral aliquot at room temperature.
48. Prepare 10x serial virus dilutions:

a) Distribute 8 1.5 ml tubes in two rows (totaling 16 tubes – 1 row for each plate).
b) Identify the tubes according to the dilution (10^-1^, 10^-2^, 10-^3^, 10^-4^, 10^-5^, 10^-6^, 10^-7^, 10^-8^).
c) Using p1000, add 1,170.0 µl of infection medium to all tubes.
d) Vortex the viral aliquot
e) Add 130 µl of the virus to the tubes labeled 10^-1^, mix thoroughly with the pipette and vortex briefly.
f) For subsequent dilutions, remove 130 µl of the newly created dilution and insert it into the next tube, as shown in figure 5b.

**Critical:** For each newly created dilution, vortex briefly. This ensures that the contents of the tubes are uniform and there is no difference in the plate reading.

g) Store these tubes with the respective dilutions in the incubator at 37°C with 5% CO_2_.
49. Discard supernatants from plates containing Vero-CCL81 cells.
50. Identify the dilutions and controls on the plates, as shown in figure 5c.
51. Distribute 100 µl infection medium in rows 11 and 12 which correspond to MOCK.
52. Distribute 100 µl of each dilution in the corresponding horizontal rows (Fig. 5d).
53. Store the plates in the incubator at 37°C with 5% CO_2_ for 72 h.

**Day 5:** reading of plates by optical microscopy and calculation of viral titer

**Timing:** 2 h

54. Remove the plate from the incubator and read the wells under an optical microscope (Figure 5e).
55. Start reading through MOCK and check if there is no contamination.
56. In infected wells, one must look carefully, including in peripheral regions, to see if there is a CPE.

**Critical:** If a particular well raises doubts as to whether or not the CPE is present, view it and compare it again with the mock.

**Critical:** Use a 96-well plate reference sheet to help identify wells that do or do not show CPE. If a CPE is present in a well, mark a “+” on the reference sheet. If there is no CPE in a well, mark a “-” on the reference sheet.

57. Repeat steps 45 through 47 for the second board.
58. After reading both plates, obtain the TCID_50_/ml titer for this stock.

**Critical:** both plates must show the same viral titer or a deviation ≤ 5%.

### Stage 5: co-infection of the A549 cell line by Mtb and SARS2

**Timing:** 5 d

**Critical step:** To carry out this step, all previous steps must have been carried out.

**Day 01:** plating of A549 cell line

59. In a BSL-1 or BSL-2 laboratory, seed A549 cells in a 12-well plate at a concentration of 5 x 10^4^ cells/well, considering 1 ml of maintenance medium per well.
60. To obtain ideal confluence, keep the plates in an incubator at 37°C with CO_2_ for 48 h.

**Dia 02:** preparation of bacterial inoculum and infection of A549 cells

**Timing:** 4 h

**Critical:** Mtb handling must be carried out exclusively in BSL-3 laboratories.

61. Transfer the plates to the BSL-3 laboratory and store them in the incubator at 37°C with 5% CO_2_ until infection is performed.
62. After 28 days of bacterial growth, check the OD in the spectrophotometer:

a) Pipette 2 ml of 7H9 Medium into a cuvette to serve as the “blank”.
b) Pipette 2 ml of the bacterial culture into another cuvette.

**Critical:** Mtb is a pathogen whose main means of transmission is the emission of aerosol particles, therefore, when leaving the biosafety cabinet and taking the sample to the spectrophotometer, place parafilm in the upper region of the cuvette containing the bacterial culture.

c) Set the spectrophotometer to measure at 600 nm (Fig. 6).
d) An OD 600 nm of 0.6 represents approximately 1×10^8^ bacteria/mL.

63. To obtain an appropriate volume of bacteria for ∼1×10^8^ bacteria in 5 mL of supplemented 7H9 medium, transfer 1 mL of the culture to a 15 mL conical tube and add 4 mL of supplemented 7H9.
64. Centrifuge for 10 min at 1,800 x g at 37°C.
65. Discard the supernatant.
66. Resuspend the bacterial pellet in 1 ml of infection medium in the 15 mL conical tube.
67. Place the tube on a 15 ml conical tube rack to provide support, and with the 1 ml syringe resuspend the bacterial solution ten times or until the bacteria are completely disaggregated.

**Figure 6.**
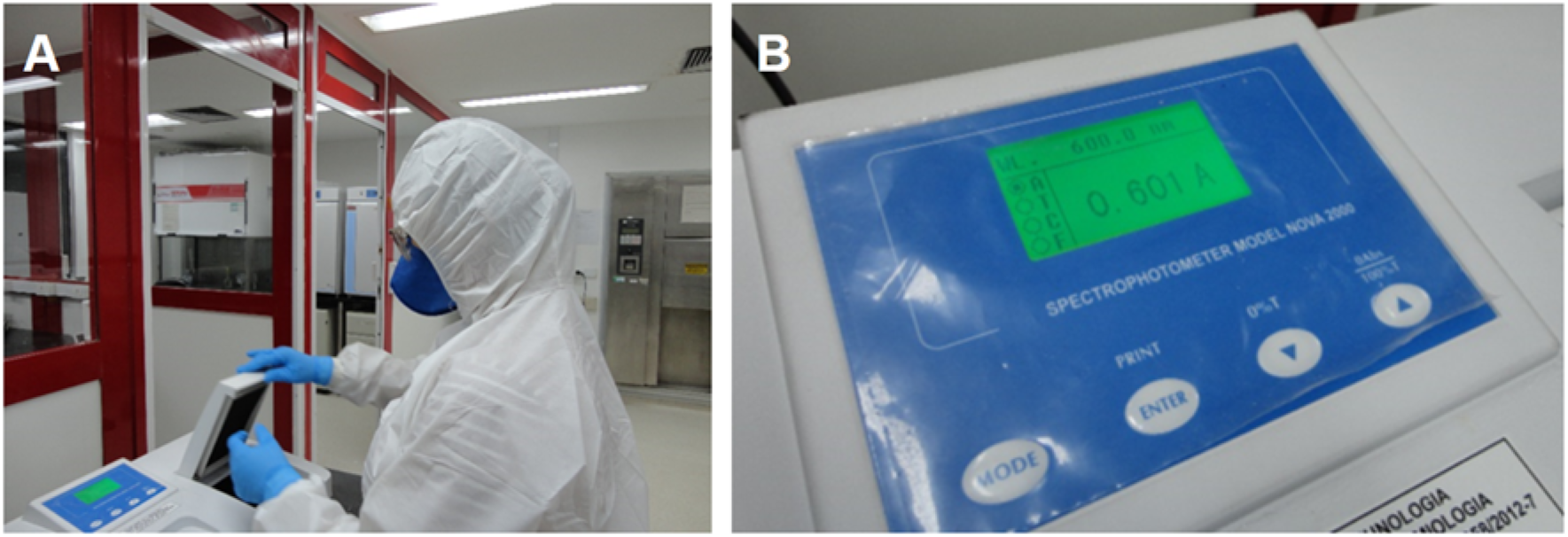
OD measurement by Spectrophotometry. A) Experimentalist measuring OD. B) Confirmation of OD at 0.6 nm.

**Critical:** *Mycobacteria* grow in clusters, which makes it difficult to break them up. If the volume of 1 ml is too low and makes it difficult for the syringe to reach the culture, add more medium, as long as it does not exceed 10 ml.

68. Add 9 mL of infection medium at a tenfold dilution (homogenize with a pipette) to approximately 1×10^7^ cells/ml.
69. Store the culture at 37°C until infection is performed.
70. Remove the culture plates from the incubator and remove the supernatant.
71. Add an MOI of 10:1 (bacteria:cell) to the wells specified for bacterial mono-infection and co-infection, corresponding to 1 ml per well.

**Critical:** For infection of A549 cells in a 12-well plate, consider the cell population of each well as 2}10^5^, adjust the concentration of bacteria to 2}10^6^ and calculate the required volume to 1 ml per well.

72. Incubate the plates for 2 h at 37°C with 5% CO_2_.
73. Carefully remove the supernatant from the wells and avoid disturbing the cell’s monolayer.
74. Wash the cells twice with 0.5 ml PBS.
75. Add 1 ml of infection medium.
76. Incubate the plates for 1 h at 37°C with 5% CO_2_.
77. Repeat steps 73 and 74.
78. Add 1 ml of infection medium and store the plates in the incubator at 37°C with 5% CO_2_ for 24 h.

**Day 03:** preparation of viral inoculum and infection of A549 cells

**Timing:** 2 h

79. Continue assuming a cell population of 2 x 105 cells/well. Dilute the viral stock (obtained in step 3) to an MOI of 1:1 (virus:cell) in a volume of 100 μl per well with infection medium.
80. Remove the plates from the incubator and in the biosafety cabinet remove the medium from the plates.
81. Wash twice with PBS at approximately 0.5 ml per well.
82. Remove the PBS and add 100 μl of the viral inoculum to the wells corresponding to SARS2 monoinfection and co-infection (Fig. 7).

**Figure 7.**
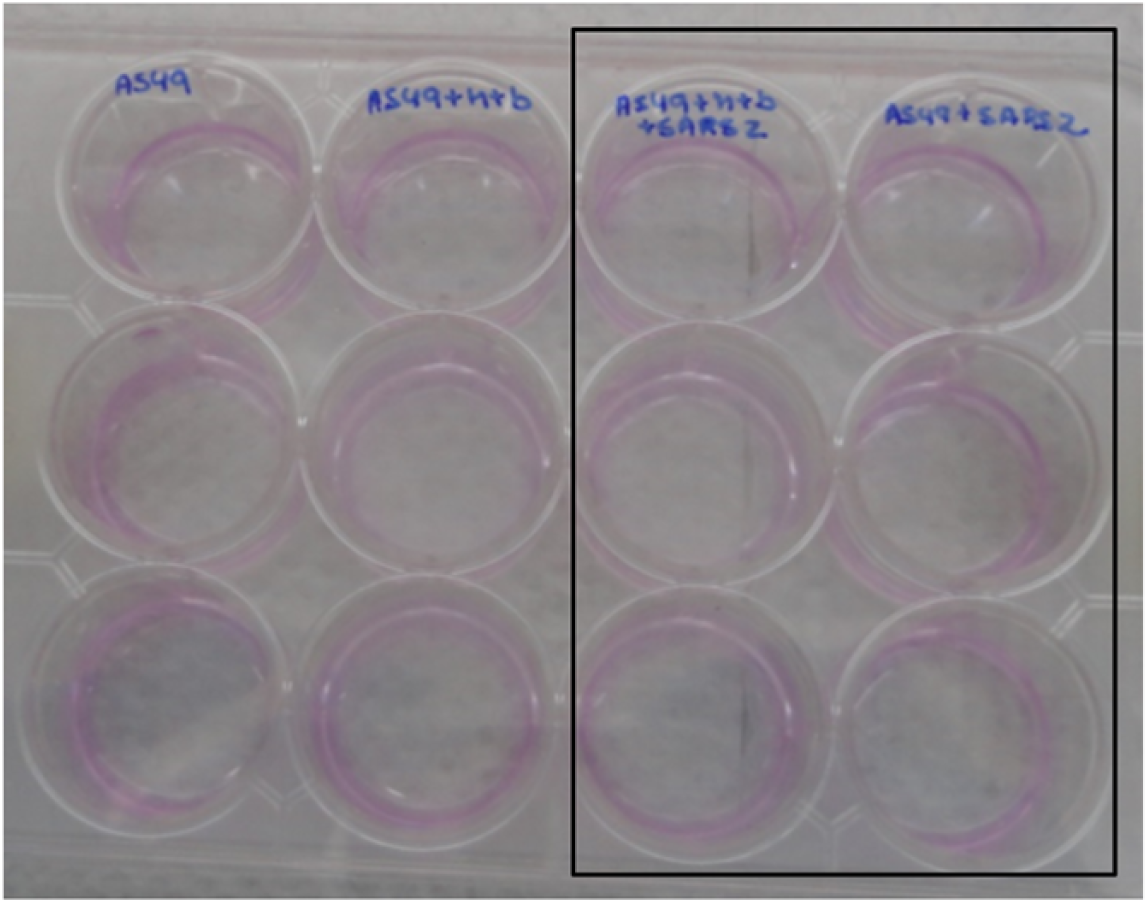
Virus inoculation into wells A549 + Mtb + SARS2 and A549 + SARS2 in.

**Critical:** it is important to have a “mock” well, where only 100 μl of infection medium but no virus will be added.

83. Return the plate to the incubator at 37°C with 5% CO_2_ for 1 h.

**Critical:** every 10 minutes, gently make circular movements on the plates. This procedure serves to redistribute the inoculum and prevent the cell monolayer from drying out.

84. Return the plates to the biosafety cabinet and remove the viral inoculum.
85. Wash the wells twice with 0.5 ml of PBS and add 1 ml of infection medium to each well.
86. Incubate the plates at 37°C with 5% CO2 for 72 h.

**Day 05:** infection harvesting

**Timing:** 3 h

87. Remove the plates from the incubator.
88. In the biosafety cabinet, collect the infection media with p1000, transfer to properly identified 1.5 ml tubes, and set aside the culture plates.

**Critical:** After collecting the supernatants, do not discard the plate. It should be reserved to continue the experiments, as there will still be viable cells at the bottom of the plate well that can be used for RNA extraction and determination of the intracellular bacterial load.

89. Centrifuge the tubes at 15,000 × g for 1 min.
90. Using a 1 ml syringe, remove the supernatants from the tubes without disturbing the pellet.

**Critical:** in the tubes containing Mtb mono-infection and co-infection samples should be stored. These cell products will be returned to their respective culture wells for Mtb intracellular bacterial load testing and RNA extraction. Further details are described in stage 6.

91. Attach the syringe to a 0.22 µm filter and filter the supernatant into a new tube.

**Critical:** Supernatants from SARS2 mono-infection and co-infection wells can be used for quantification of viral particles by TCID50 and protein analysis by flow cytometry. To do this, it is necessary to aliquot these supernatants into different appropriately labeled tubes. The samples obtained can be measured immediately or stored at -80°C until needed.

92. Store all supernatants at -80°C.
93. Return the pellet to its respective well and perform RNA extraction using the Trizol method or proceed to the determination of intracellular bacterial load (Stage 6, steps 94-101).

### Stage 6: determination of intracellular bacterial load

**Timing:** 2 h

94. In the reserved plate, return the pellets obtained after centrifugation to the respective wells (step 5, step 90).
95. Using p1000, add 1 ml of lysis buffer to the wells corresponding to Mtb mono-infection and co-infection (Fig. 8a).
96. Incubate the plate at room temperature for 10 min to promote cell lysis (Fig. 8b).
97. Observe the cells under an optical microscope; you will be able to see the Mtb-infected cells “swollen” (Fig. 9).

**Figure 8.**
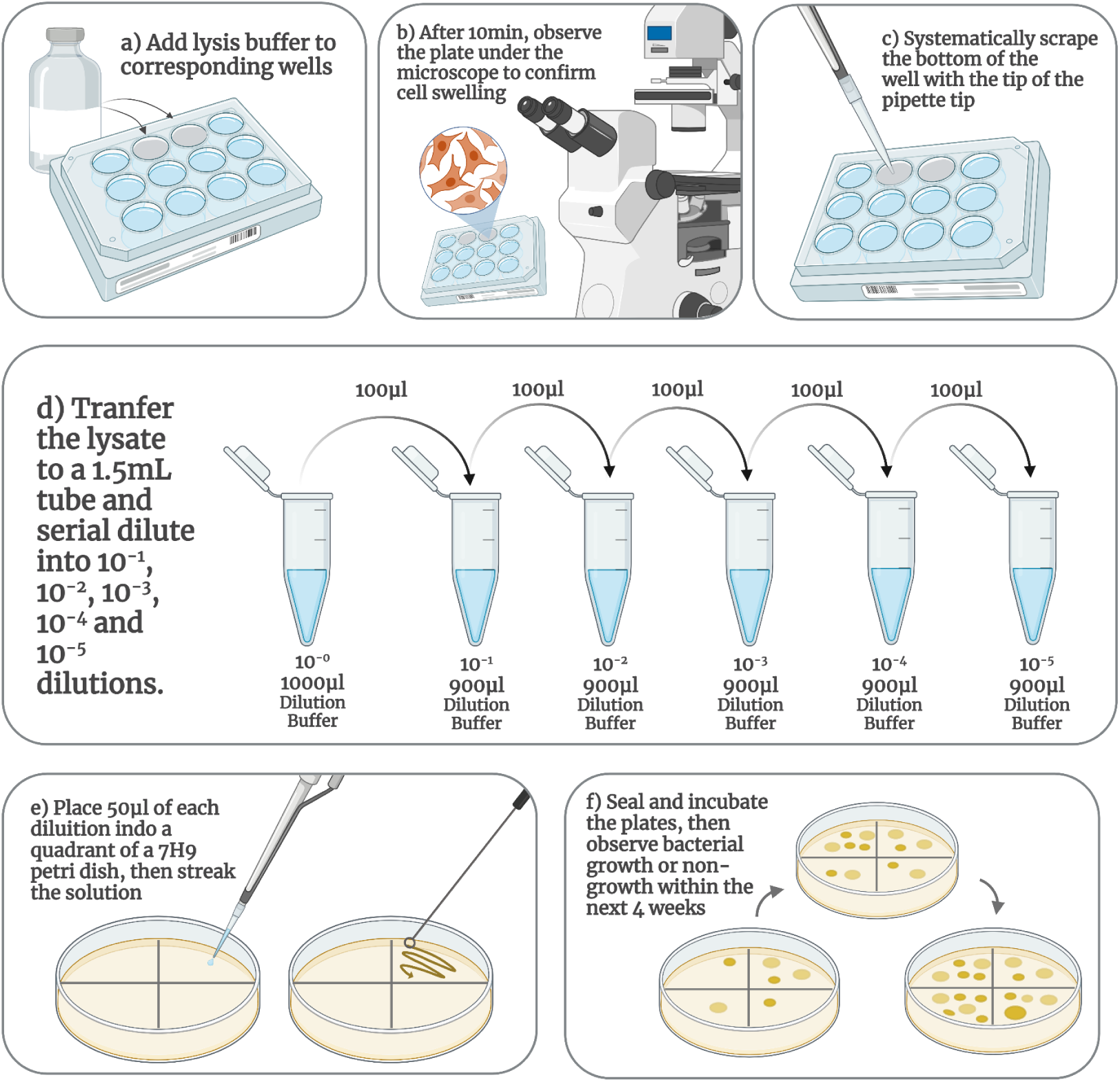
Workflow for determination of intracellular bacterial load. a) Lysis buffer in the specific well. b) Observing the activity of the lysis buffer under the microscope. c) Technical scraping to obtain better lysates. d) Serial dilution of lysates. e) Plating of dilutions. f) CFUs count.

**Figure 9.**
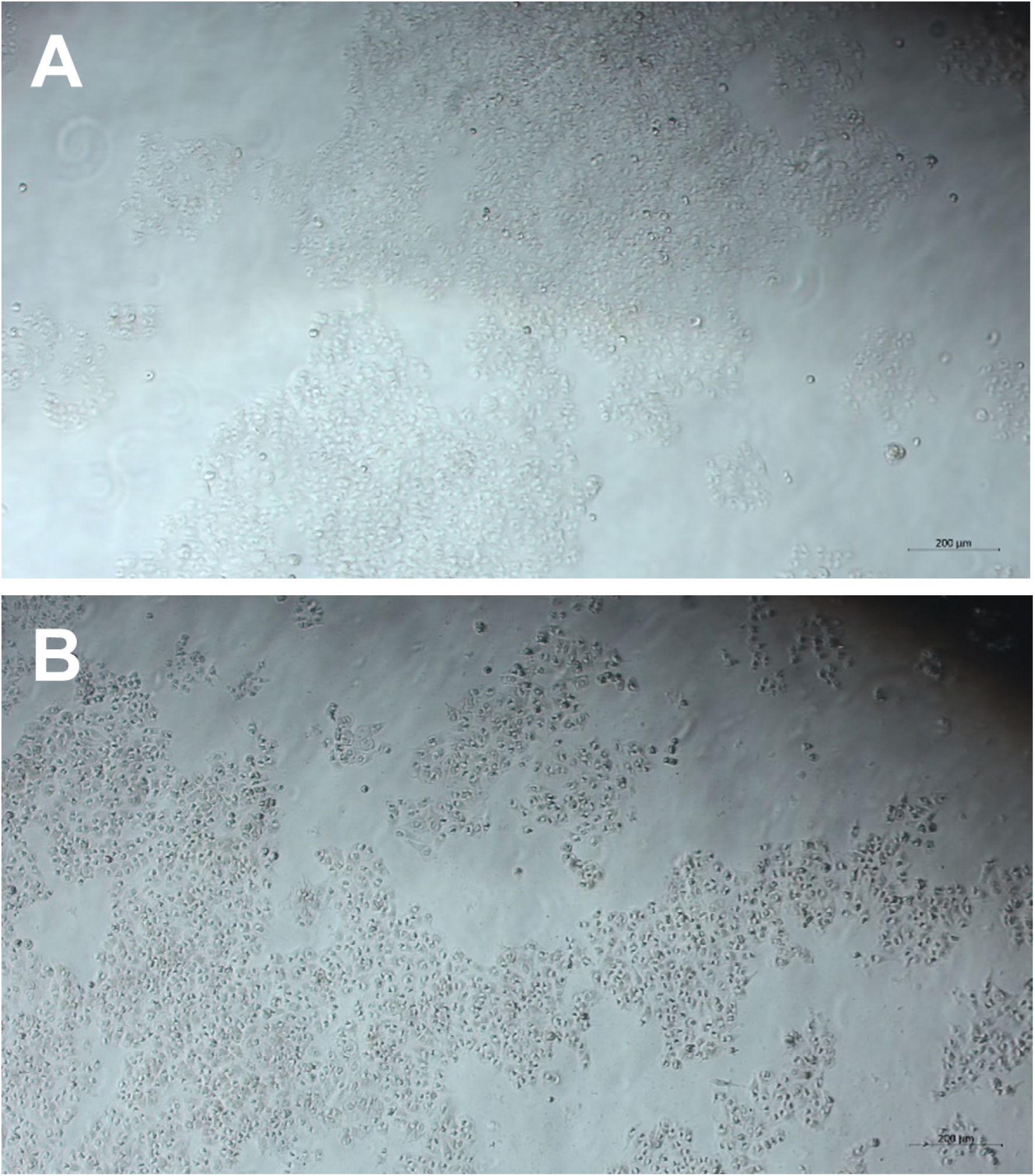
Mtb infection and wells with lysis buffer. A) A549 + Mtb and with lysis buffer. B) A549 + Mtb + SARS2 and with a lysis buffer.

**Critical:** note the difference between uninfected cells and cells with lysis buffer.

98. Using a p1000, systematically scrape the well with the tip of the tip (Fig. 8c), and pipette the suspension equally into all wells up and down. This technique will promote complete and linear lysis of the cells.
99. In 1.5 ml tubes, serially dilute the lysed cells in suspension by 10^−1^, 10^−2^, 10^−3^, 10^−4^ and 10^−5^. Follow the image in figure 8d to perform the dilutions.

**Critical:** save all unplated tubes diluted and undiluted until the end of the CFU count.

100. Place 50 μL of each dilution on a quadrant of the appropriate 7H10 agar plate. Spread the lysed cell solution onto the sterile 7H10 plate with a disposable inoculating loop and allow it to dry completely (Fig. 8e).
101. Seal the plates with parafilm and incubate at 37°C. Colony forming units (CFUs) will begin to appear approximately 2 to 3 weeks after inoculation (Fig. 8f).

**Critical:** perform a 7H10 control plate with the reagents used in the bacterial culture to ensure the absence of contamination.

**Critical:** it is important that the plates are checked after 48 hours of inoculation to ensure the absence of contamination.

**Critical:** when colonies begin to appear, they should be counted weekly to check whether there has been more growth in CFUs from one week to the next. Plates with 7H10 should be stored for up to 28 days. After this period, they should be discarded.

### Stage 7: viability test of SARS2 by TCID_50_ assay

**Timing:** 5 d

### Day 1: plating of Vero-CCL81 cells for infection

**Timing:** 1 h plating and overnight incubation

102. Plating of Vero-CCL81 cells (Fig. 10a) for this step can be performed as per stage 4, step: 35 and 36.

**Figure 10.**
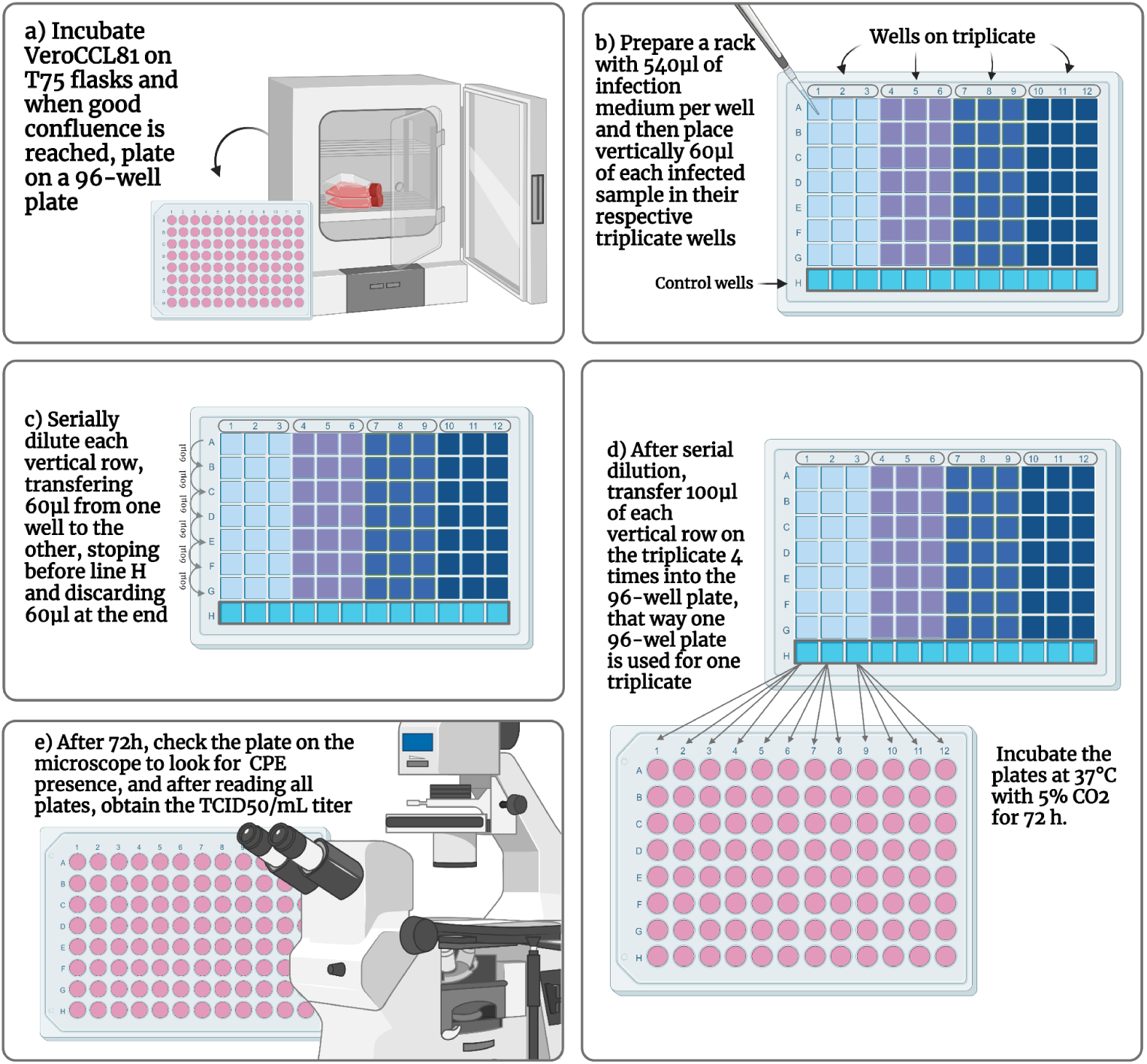
Workflow of SARS2 viability test. a) Vero-CCL81 incubation. b) Preparation of the rack with the supernatants and infection medium. c) Serial dilution in the rack. d) Every third vertical row of the rack, transfer to a 96-well plate. e) Observation of CPE under the microscope.

**Critical:** calculate the number of possible 96-well plates according to the number of samples that need to be quantified. Each triplicate of samples is equivalent to a 96-well plate. Each rack is equivalent to 4 triplicates of samples (Fig. 10b), resulting in a total of 4 96-well plates.

### Day 2: rack preparation and infection of Vero-CCL81 cells with supernatants

**Timing:** 1 h rack preparation, 1.5 h infection, 72 h incubation

103. Forward the 96-well plates to the BSL-3 laboratory.
104. In a biosafety cabinet, distribute 540 µl of infection medium into each well of the rack (Fig. 10b).
105. Add 60 µl of each sample to the first well of each vertical row.
106. Perform serial dilution of each vertical row, transferring 60 µl from one well to the other (Fig. 10c).

**Critical:** in well G of each vertical row, add 60 µl and then discard the sample. Do not pipette anything into well H of each vertical row because it is the control.

107. Transfer 100 µl from the rack to the 96-well plate (Fig. 10d).

**Critical:** each vertical row will be added 4x in the 96-well plate (also vertically), where each triplicate (obtained from cell culture) of samples will be a 96-well plate.

108. Incubate the plates at 37°C with 5% CO_2_ for 72 h.

**Day 5:** reading of plates by optical microscopy and calculation of viral titer

**Timing:** 3 h

109. Today’s procedure can be performed as per stage 4, step: 54-58, and both can be viewed as per figure 11.

**Figure 11.**
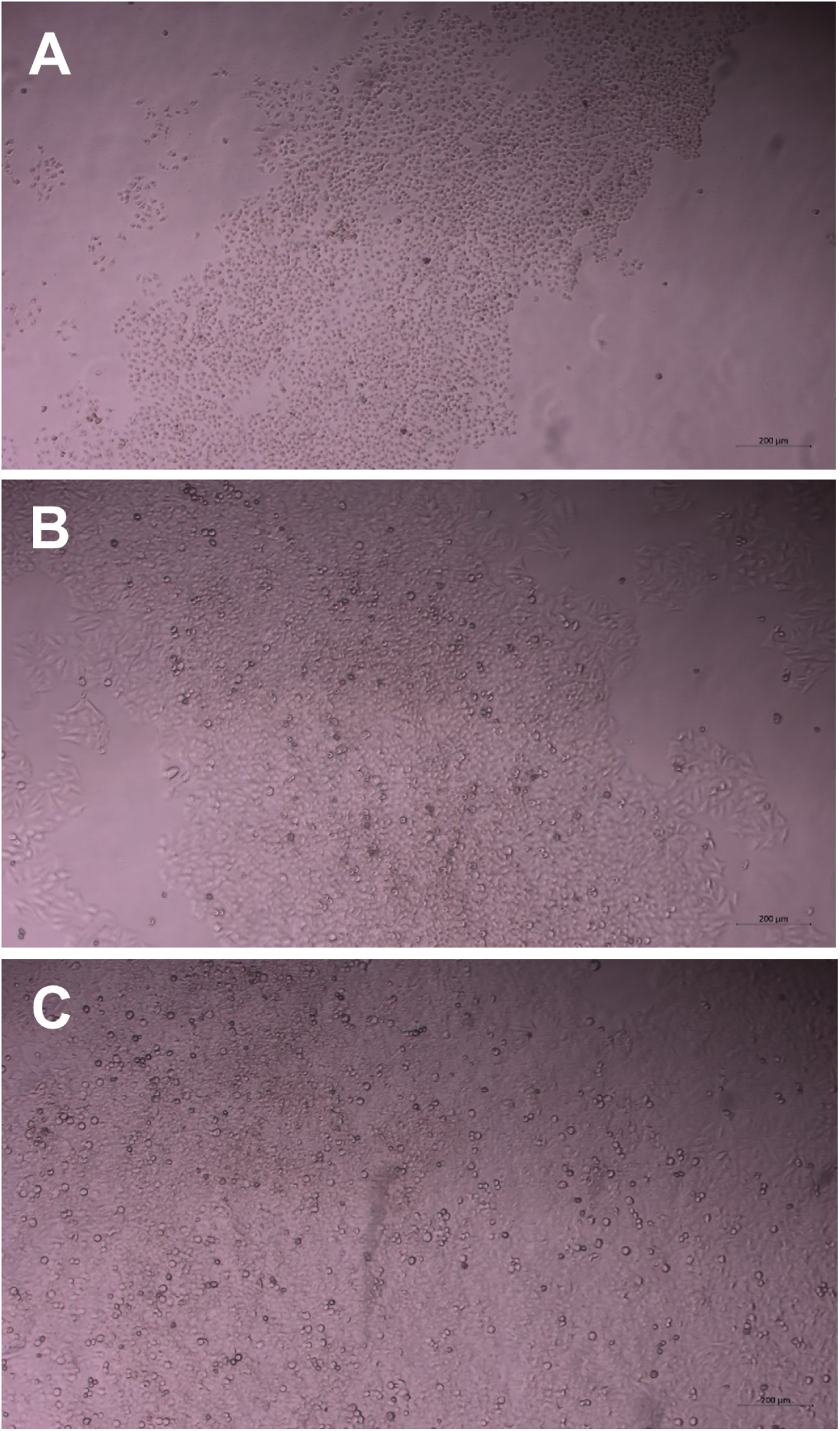
TCID_50_ assay and CPE visualization. A) Control well (A549 only). B) A549 + Mtb + SARS2. C) A549 + SARS2.

### Stage 8 (optional): decontamination of supernatants by UV-C

**Timing:** 5 d

**Day 1:** preparation of Vero-CCL81 cells

**Timing:** 1 h plating, overnight incubation

**Critical:** this process can be performed in BSL-1 or BSL-2.

110. In a 6-well culture plate, plate a concentration of Vero CCL-81 cells corresponding to 2 x 10^5^ cells/ml in each well (Fig. 12a).
111. Incubate the plate overnight in an incubator at 37 °C with 5% CO_2_.

**Figure 12.**
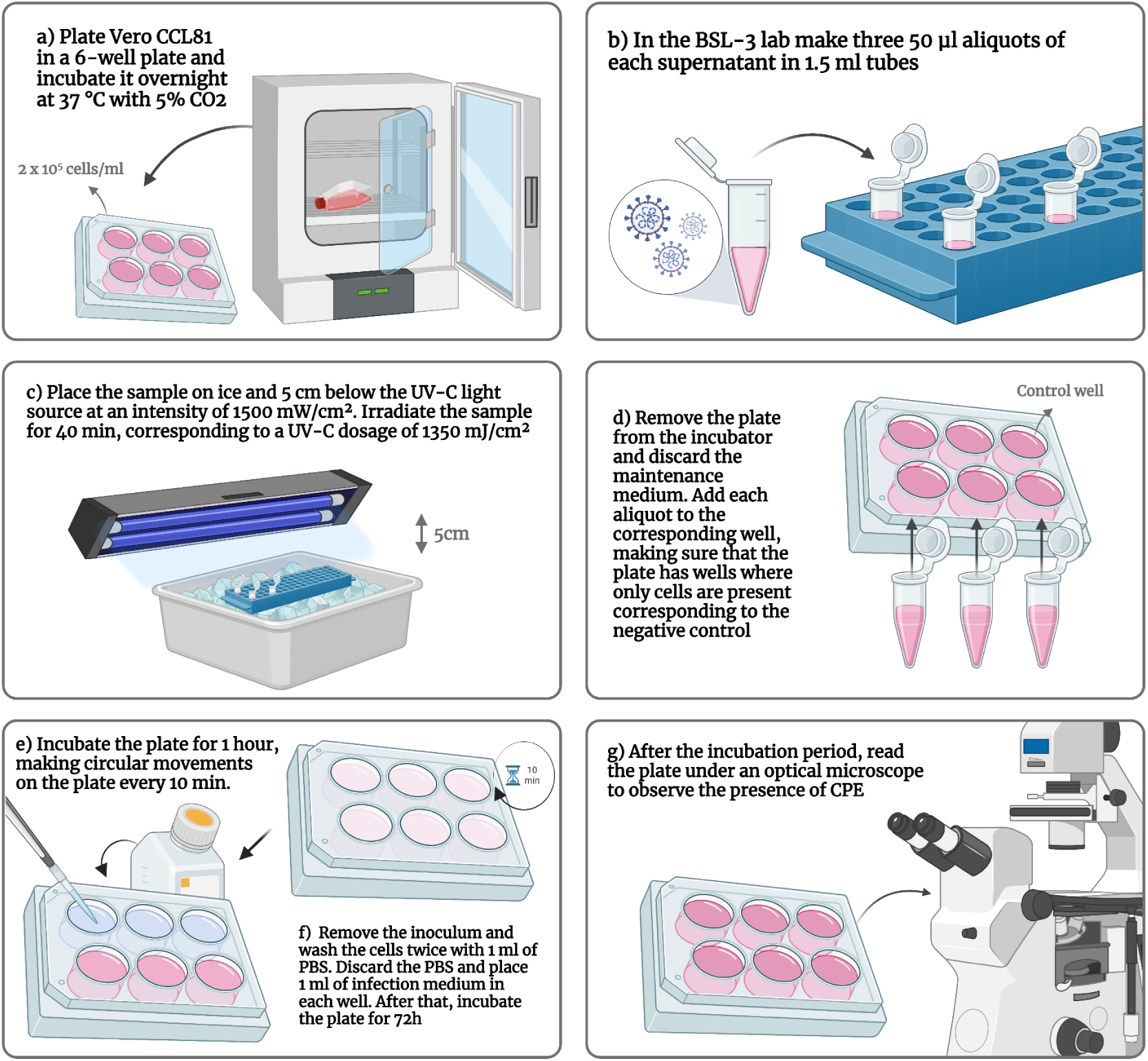
Workflow for supernatant decontamination. a) Vero CCL-81 incubation. b) Aliquots of the supernatant. c) UV-C radiation at 5 cm for 40 min. d) Preparation of the 6-well plate with the viral aliquots and negative control wells (cells only). e) Incubation of the aliquots for 1 h and circular movements every 10 min. f) Washing the wells and adding infection medium. g) Incubation for 72 h and observation of CPE under the microscope.

**Day 2:** sample decontamination test **Timing:** 2 h decontamination, 72 h incubation. **Critical:** this process must be performed in BSL-3.

112. After the incubation period, transfer the plate to BSL-3.
113. Make 3 aliquots of 50 µl of each supernatant in 1.5 ml tubes (Fig. 12b).

**Critical:** the 3 aliquots must come from the same supernatant, and each one must be inoculated into three different wells.

114. Place the sample on ice and 5 cm below the UV-C light source at an intensity of 1500 mW/cm² as per figure 12c;
115. Irradiate the sample for 40 min, corresponding to a UV-C dosage of 1350 mJ/cm².

**Critical:** It is necessary to check how long it takes for the UV-C light source used to reach this intensity and dosage, and then count the 40 minutes.

116. Remove the plate from the incubator and discard the maintenance medium;
117. Add each aliquot to the corresponding well for the decontamination test (Fig. 12d).

**Critical:** make sure that the plate has wells corresponding to the negative control, where there will only be cells, without insertion of the sample that will be decontaminated.

118. Incubate in the incubator for 1 hour, and every 10 min make circular movements on the plate (Fig. 12e).
119. Remove the inoculum and wash the cells twice with 1 ml of PBS (Fig. 12f).
120. Discard the PBS and place 1 ml of infection medium in each well. Incubate the plate in an incubator at 37°C with 5% CO_2_ for 72 h.

**Day 5:** reading the plate by optical microscopy

**Timing:** 1 h

121. After the incubation period, read the plate under an optical microscope to observe the CPE (Fig. 12g).

**Critical:** the absence of CPE in the wells corresponding to viral aliquots demonstrates efficient sample decontamination.

### TIMING

Stage 1, Mtb culture: 28 d after thawing

Steps 1-3, preparation of bacterial culture: 1 h
Steps 4-6, bacterial culture maintenance: 1 h, optimal growth culture: up to 28 d.

Stage 2, cell maintenance: 10 d after thawing

Steps 7-17, thawing of cells: 1 h, replating and counting for growth: 2-3 d.
Steps 18-28, replating of cells: 1 h, cell division: 4-6 d for two passages
Stage 3, preparation of viral stocks: 5 d.
Steps 29-30, plating of Vero-CCL81 cells: 1 h.

Steps 31-36, infection of Vero-CCL81 cells with the hCoV-19/Brazil/PE-FIOCRUZ-IAM4372/2021 strain: inoculation 2 h, incubation 72 h.

Steps 37-43, virus harvest: 2 h.

Stage 4, determination of infectious titers of viral stocks: 5 d.

Steps 44-45, plating of Vero CCL81 cells: 1 h, incubation 12-16 h.
Steps 46-53, TCID_50_ assay: inoculation 2.5 h, incubation 72 h.
Steps 54-58, reading of plates by light microscopy and calculation of viral titer: 2 h.

Stage 5, co-infection of A549 cells: 5 d.

Steps 59-60, plating of A549 cells: plating 1 h, incubation 12-16 h.
Steps 61-78, preparation of bacterial inoculum and infection of A549 cells: 4 h.
Steps 79-86, preparation of viral inoculum, infection and co-infection of A549 cells: infection 3 h, incubation 72 h.
Steps 87-92, harvest of RNA and proteins: 4 h.

Stage 6, determination of intracellular bacterial load: 28 d.

Steps 93-100, cell lysis, dilutions and inoculation of plates 7H10: preparation of viability 3 h, incubation 28 d.

Stage 7, SARS2 viability test by TCID50: 5 d.

Step 101, plating of Vero CCL-81 cells: 1 h plating, 12-16 h incubation.
Steps 102-107, rack preparation and inoculation with viral harvests (mono- and co-infected) from the culture: 3.5 h cultivation, 72 h incubation.
Step 108, reading of plates by light microscopy and calculation of viral titer: 3 h.

Stage 8, UV-C decontamination: 5 d.

Steps 109-110, plating of Vero CCL-81 cells: 1 h plating, 12-16 h incubation.
Steps 111-120, preparation of supernatant aliquots: 2 h decontamination, 72 h incubation.
Step 121, reading of plates by light microscopy: 1 h.

## ANTICIPATED RESULTS

### Co-infection Mtb + SARS2

After successful cultivation of Mtb and SARS2 individually, we performed mono- and co-infections in the host cell, and after 72 h of completed infections we visualized the effects on the cells via microscopy. CPE in A549 cells used in this protocol will present as cell death resulting in cell rounding and detachment. CPE induced in mono- and co-infections can be distinguished due to super confluence of cells and therefore comparison with uninoculated control cells plated at the same time is critical.

### Determination of intracellular bacterial load

After mono-infection with Mtb and Mtb + SARS2 co-infection, we hope to detect the intracellular bacterial load through CFUs to confirm the viability of the system, and to determine the rate of infection and spread between Mtb and Mtb + SARS2. The amount of bacterial load may vary due to the dilutions used in the inoculation of the 7H10 agar medium. This load determination may also fail in some cases due to the preparation of the plates with 7H10 medium or due to the low amount of bacteria used. To ensure the success rate of infection, we recommend using the MOI of 10 for each well of infected cells.

### Determination of the infectious titer by SARS2

In mono-infection with SARS2 and Mtb + SARS2 co-infection, we expect the presence of CPE to be observed in order to, as well as the determination of the bacterial load, confirm the viability of the system, and verify the rate of infection and spread in SARS2 and Mtb + SARS2.

### Obtaining cell extracts

We hope to obtain RNA and protein samples with a high level of purity, quality and efficiency, providing experimentalists with materials to be used in other activities and to obtain other results, according to the objectives.

### UV-C decontamination

UV-C decontamination causes a rupture of the genetic material of Mtb and SARS2, making the pathogens non-infectious, but preserving the protein architecture of the cells. This decontamination will allow experimentalists to remove the material from BSL-3 to BSL-2 and continue with other experiments, according to the desired objectives.

### Abstract

This protocol efficiently produces a co-infection system between two distinct pathogens. Mtb and SARS2, alone or together, can cause CPE in A549 cells. In this system, it is necessary to confirm the viability of each of the pathogens, which allows determining the rate of infection and spread between mono-infections: Mtb and SARS2, and co-infection: Mtb + SARS2. Highly pure RNA and protein samples can be obtained from the cultures, which can undergo a decontamination step, providing experimentalists with a safe exit from BSL-3 to BSL-2.

## TROUBLESHOOTING

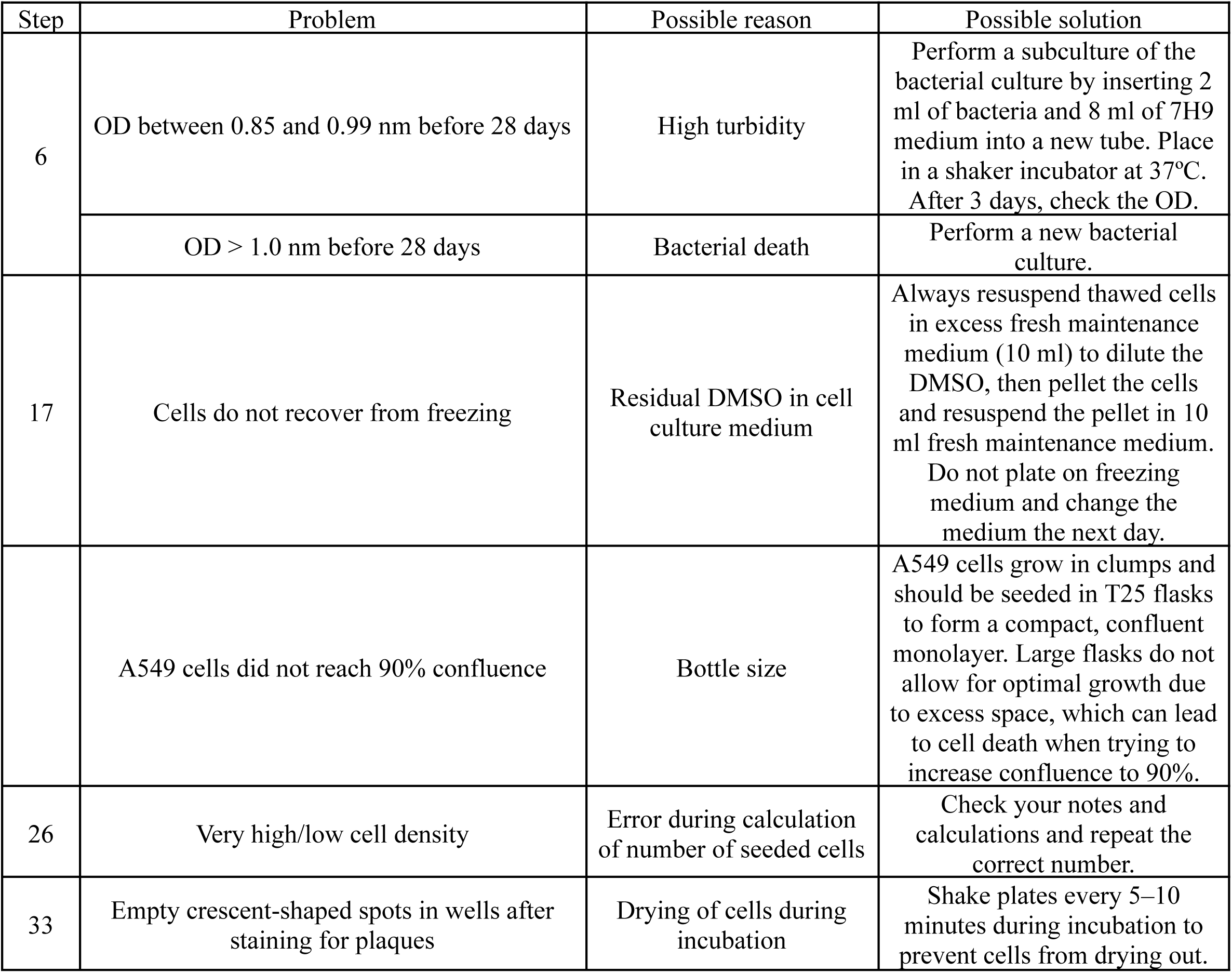
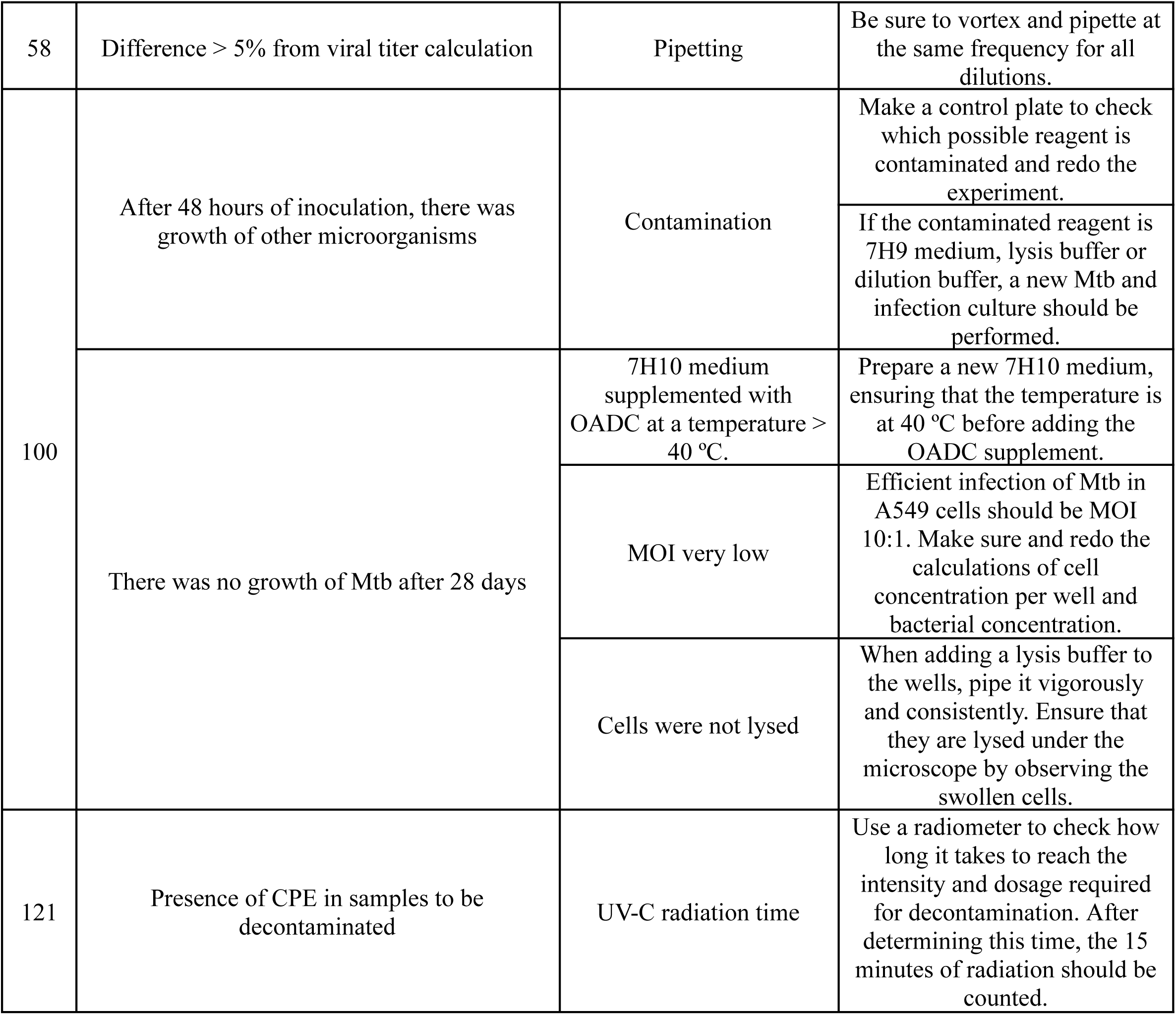

## ACKNOWLEDGMENTS

This work was supported by the Brazilian funding agencies: Conselho Nacional de Desenvolvimento Científico e Tecnológico (CNPq) - Process Number: 140437/2022-3; Universal - CNPq - Process Number: 403844/2021-5; Fundação de Amparo a Ciência e Tecnologia de Pernambuco (FACEPE) - Process Number: APQ-1119-2.02/22.

## AUTHOR CONTRIBUTIONS

Thays Maria Costa de Lucena: Writing – original draft & editing, Methodology, Formal analysis, Conceptualization. Débora Elienai de Oliveira Miranda: Methodology. Juliana Vieira de Barros Arcoverde: Methodology. Mariana Souza Bezerra Cavalcanti: Methodology. Michelle Christiane da Silva Rabello: Methodology. Jaqueline de Azevêdo Silva: Funding acquisition, Writing – review & editing, Validation, Supervision, Conceptualization.

## REFERENCES

1. Global Tuberculosis Report 2023. (2023).

2. Khurana, A. K. & Aggarwal, D. The (in)significance of TB and COVID-19 co-infection. European Respiratory Journal 56, 2002105 (2020).

3. Crisan-Dabija, R. et al. Tuberculosis and COVID-19: Lessons from the Past Viral Outbreaks and Possible Future Outcomes. Can Respir J 2020, 1–10 (2020).

4. Gao, Y. et al. Association between tuberculosis and COVID-19 severity and mortality: A rapid systematic review and meta-analysis. J Med Virol 93, 194–196 (2021).

5. Sy, K. T. L., Haw, N. J. L. & Uy, J. Previous and active tuberculosis increases risk of death and prolongs recovery in patients with COVID-19. Infect Dis 52, 902–907 (2020).

6. Dheda, K. et al. The intersecting pandemics of tuberculosis and COVID-19: population-level and patient-level impact, clinical presentation, and corrective interventions. Lancet Respir Med 10, 603–622 (2022).

7. Chiok, K. R., Dhar, N. & Banerjee, A. Mycobacterium tuberculosis and SARS-CoV-2 co-infections: The knowns and unknowns. iScience 26, 106629 (2023).

8. Udwadia, Z. F. et al. COVID-19 -Tuberculosis interactions: When dark forces collide. Indian Journal of Tuberculosis 67, S155–S162 (2020).

9. Coronavirus disease (COVID-19). https://www.who.int/emergencies/diseases/novel-coronavirus-2019/advice-for-public?adgroupsurvey={adgroupsurvey}&gad_source=1&gclid=EAIaIQobChMIk8Cbk6XlhwMVXWBIAB3MBAjOEAAYASAAEgLKCfD_BwE.

10. Goncheva, M. I. & Heinrichs, D. E. Protocol for studying co-infection between SARS-CoV-2 and Staphylococcus aureus in vitro. STAR Protoc 4, 102411 (2023).

11. Rosli, S. N. Z. et al. Vero CCL-81 and Calu-3 Cell Lines as Alternative Hosts for Isolation and Propagation of SARS-CoV-2 Isolated in Malaysia. Biomedicines 11, 1658 (2023).

